# Basal Ganglia Responses to Electrical Stimulation of the Posterior Hypothalamic Nucleus

**DOI:** 10.1101/2021.05.17.444549

**Authors:** Calvin K. Young, Brian H. Bland

## Abstract

Electrical or chemical stimulation of the posterior hypothalamic nucleus (PH) elicits highly adaptive locomotion, demonstrating both evidence of flexibility and variety in exhibited motor behaviours. However, the neural substrates of PH stimulation elicited behavioural changes are poorly understood. The basal ganglia are postulated to be critically involved in the process of action selection in conjunction with thalamo-cortical systems. The present study examines changes in basal ganglia activities in response to the high-frequency stimulation of the PH. Under urethane anaesthesia, ensemble and single-unit recordings were obtained from the striatum (STR), globus pallidus externa (GPe), entopeduncular nucleus (EP), subthalamic nucleus (STN) and the substantia nigra pars reticulata (SNr). Upon PH stimulation, increases in firing rates were observed in the STR, GPe, and STN, a decrease was observed in the SNr and no changes were seen in the EP. The increase in spike rate in the STR and GPe was dependent on the stimulation intensity but not duration. Despite the differences in the direction of firing changes during PH stimulation, all examined areas including those not part of the basal ganglia demonstrated an elevated spiking rate upon stimulus train termination. Taking into account the known anatomical connections between the PH and the basal ganglia, it is hypothesized responses seen during PH stimulus trains are mediated through thalamic and cortical relays whereas the overall post-stimulus excitatory response is related to the impact of the PH on brainstem arousal systems.

---

The basal ganglia are a collection of highly interconnected subcortical nuclei that receive divergent thalamic and cortical inputs (Mengual et al., 1999; Van der Werf et al., 2002) (van der Kooy, 1979; Veening et al., 1980; Fuller et al., 1987; Berendse and Groenewegen, 1990; Sadikot et al., 1992; Erro et al., 2001; Erro et al., 2002). Through a series of inhibitory controls within the basal ganglia, the information entering the striatum (STR) either feeds back to the motor thalamus through the internal segment of the globus pallidus (GPi, or the entopeduncular nucleus in rodents; EP) or travels through the globus pallidus externa (GPe) to modulate the excitatory subthalamic nucleus (STN) control over the substantia nigra pars reticulate (SNr), which also projects to the motor thalamus. Direct cortico-STN connections (Carpenter et al., 1981; Kitai and Deniau, 1981; Fujimoto and Kita, 1993; Nambu et al., 2002), direct STR-SNr projections (Parent and Hazrati, 1995a), and all the intricate details in the topographical organization of projection patterns between different nuclei in the basal ganglia adds to the complexity of the information processing pathways through the circuitry.

The posterior hypothalamic nucleus (PH) is a large midline hypothalamic nucleus that has a prominent role in the modulating changes of cardiovascular and motor functions (Bland and Vanderwolf, 1972; Shekhar and DiMicco, 1987). It has long been demonstrated that chemical or electrical stimulation of the PH or surrounding areas results in the elicitation of locomotion (Bland and Vanderwolf, 1972; Shekhar and DiMicco, 1987; Lammers et al., 1988; Marciello and Sinnamon, 1990; Sinnamon et al., 1991; Slawinska and Kasicki, 1995; Oddie et al., 1996; Slawinska and Kasicki, 1998). The behavioural characteristics of elicited locomotion have been described to resemble either exploratory behaviour or defensive flight, which may be dependent on the intensity of the stimulation (Shekhar and DiMicco, 1987; Lammers et al., 1988). Regardless of its nature, the elicited locomotion is described as non-stereotypical, meaning a larger repertoire of motor behaviours are expressed, such as running, jumping, climbing and rearing, depending on the physical environment the animal is in (Bland and Vanderwolf, 1972; Jackson et al., 2008). The variety of highly integrated motor behaviours is suggestive of involvement from higher-order motor areas, such as the motor cortex and thalamus, the cerebellum, and the basal ganglia. Incidentally, the PH provides projections to all the above areas, of which the motor cortex and the cerebellum are reciprocally connected to the PH (Vertes et al., 1995; Vertes and Crane, 1996; Abrahamson and Moore, 2001; Cavdar et al., 2001b; Cavdar et al., 2001a). Previous studies have revealed an excitatory effect of PH stimulation on the cortex (Young et al., 2011; Young and Bland, 2021). Particularly, it was shown that motor cortical spiking evoked by PH stimulation is blocked by motor thalamic inactivation, suggesting PH stimulation excites thalamic neurons which in turn excites cortical neurons. Under anaesthesia, PH stimulation was also able to reduce the current threshold of intracortical stimulation-induced movements.

While the relationship of PH-thalamo-cortical axis has been examined, the effects of PH stimulation on other higher-order motor areas remain unknown. The basal ganglia are postulated to integrate divergent sensorimotor information from all over the cortex to provide instructions to the motor cortex via the motor thalamus, among other systems (Graybiel, 1995; Mink, 1996; Graybiel, 1998; Redgrave et al., 1999; Grahn et al., 2008; Seger, 2008). The flexibility of motor behaviours displayed in PH-stimulated rats suggests basal ganglia involvement. In this study, spiking responses from the STR, GPe, EP, STN, and SNr were examined with different PH stimulation protocols. The PH only provides sparse and light innervations to the STR, GPe and the dopaminergic substantia nigra pars compacta within the main basal ganglia nuclei. Therefore, it is likely that the most consistent responses elicited from the basal ganglia nuclei would be mediated through the cortex.

## Materials and Methods

### Recording Procedures

Sixteen male Long Evans hooded rats between 250-370 g were obtained from the Animal Care Facility at the University of Calgary. Rats were caged in groups of 3-4 and were given free access to food and water in a temperature, air, and light-controlled room. The rats were prepared for recording by firstly being anaesthetised by halothane (1.5% oxygen alveolar concentration), and then maintained and supplemented by ethyl carbamate (1.2 g/kg) through a jugular cannula implanted during halothane anaesthesia. Once the rats had reached a stable anaesthetic state, characterized by the slow, stationary respiration rate for at least 20 min, they were head-fixed in a stereotaxic frame for unit recordings. The rats were kept at a constant 37°C by a temperature control unit with intermittent saline (0.9% NaCl) injections in between recording sessions. All procedures used in this study strictly adhered to the guidelines of the Canadian Council on animal care and were approved by the Animal Care Committees of the University of Calgary and the National Research Council.

Subcutaneous lidocaine (Rafter 8, Calgary, AB) was administered prior to an incision along the midline to expose the skull and its suture lines. Connective tissue was pushed aside and the skull was cleaned and dried before holes were made for electrode insertion. Generally, three to five recording sites were attempted on the side ipsilateral to PH stimulating electrode placement. In some experiments, bilateral recordings were made instead. All the coordinates used for recording and stimulating are summarized in Table 1. Unit recordings were made with either glass electrodes filled with 0.5M sodium acetate with 2% Pontamine sky blue (2-8 MΩ) or platinum/iridium tetrodes (Thomas Recording Gmbh, Germany). Stimulating electrodes were made from twisted pairs of stainless steel wires with ~0.5 mm tip separation. Single stainless steel wires were used for reference and cortical LFP recordings. The recordings were fed to a 16-channel digital amplifier (A-M Systems, Carlsborg, WA) through preamplifiers, filtered between 1-300k Hz at 1000x, digitized at 250 kHz, and acquired with SciWorks (DataWave systems, Berthoud, CO).

**Table 1.**
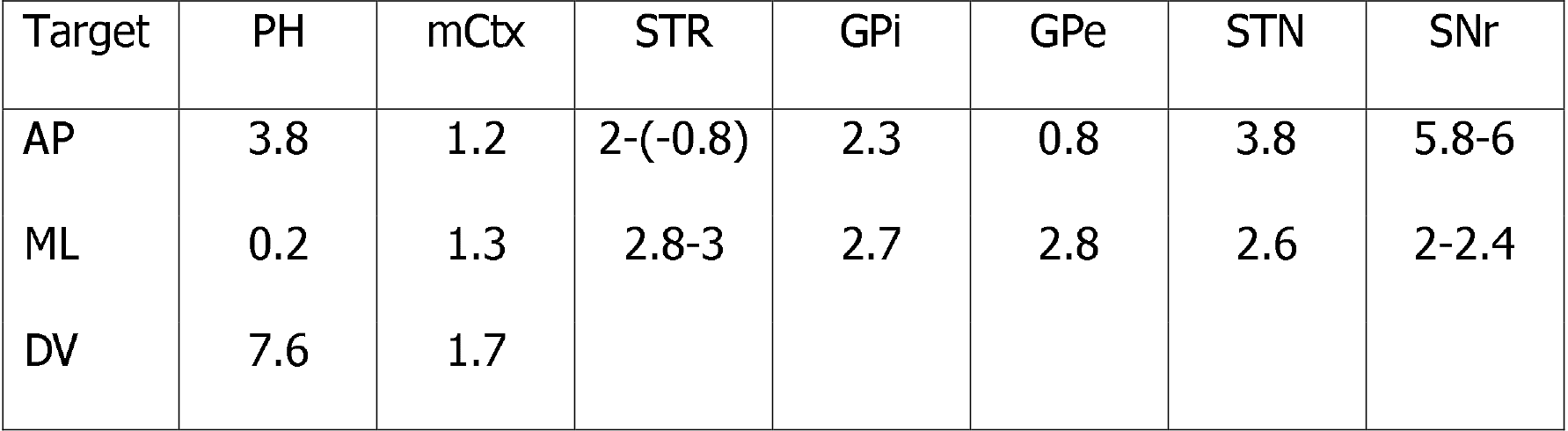
Stereotaxic coordinates for recording and stimulation.

The stimulating electrode, cortical LFP electrode, and reference electrodes were all implanted prior to lowering of recording electrodes for single or multi-unit recordings. All recording electrodes were positioned at given coordinates and lowered by using manual microdrives. During lowering of the recording electrodes, filter settings were set to .5-300 kHz bandpass, and once an acceptable signal-to-noise ratio was obtained, the electrode was left in place for at least 5 min. Since PH stimulation induces respiratory changes, once all recording sites yielded acceptable recordings, PH stimulus trains were delivered to ensure adequate recording stability and current needed to elicit cortical desynchronization. At least another 5 min was given after the last PH stimulus train before experiments began.

### Experimental Protocol

The experimental protocol used here was the same as reported previously (Young and Bland, 2021). Briefly, 2-3 min baseline recordings were made prior to the delivery of a ~10 s PH stimulus train at the intensity that evoked cortical desynchronization (0.3-0.5 mA at 100 Hz, 0.1 ms pulse duration). After the LFP recording had returned to stable UP-DOWN state transitions, single pulses (0.2 ms, 0.4 mA pulses at 0.4 Hz) were repeated at least 20 times, before the intensity and duration step protocols were administered after a brief (1-2 min) delay. Intensity step protocol (8 successive 250 ms, 100 Hz trains of 0.2 ms pulses from 0.1-0.8 mA in 0.1 mA steps) was followed by duration step protocol (9 successive 100 Hz, 0.2 ms pulses at 0.3-0.5 mA were delivered for 100-500 ms in 50 ms steps). The recording was continued for up to another 3 min to ensure spiking from recorded units was still present with the actual extended recording time depending on the baseline firing rate of the unit.

### Data Analysis

All data were imported to Matlab 7.0 (Mathworks, Natwick, MA) for analysis. Spiking data were isolated from stimulus artifacts by conventional spike-sorting techniques involving template matching and subsequent cluster analysis based on spike waveform measurements. Stimulus artifacts also had amplitudes that were magnitudes higher than action potentials, which were exploited to provide markers for triggering peri-stimulus time histogram-(PSTH) based analyses.

Baseline firing rates were obtained from averaged data from the initial 2 min recordings at the beginning of each experiment. Spike rasters were constructed by summing all spikes from each record in 50 ms bins with the total number of spikes within a bin mapped to grayscale for presentation purposes. Firing rate changes during 10 s PH stimulation were sampled by PSTHs triggered at the first and last stimulus of the 10 s train, with 3 s data before and after the trigger included for the analyses. The changes in firing rate were compared between the 3 s baseline and the 3 s immediately following PH stimulation, and between the 3 s baseline and the 3 s immediately following the termination of PH stimulation.

The analyses of responses to intensity and duration step protocols were all taken from the period of the stimulation; that is, the 250 ms during each of the intensity steps and variable duration for duration steps. The average number of spikes from the stimulation period from each unit was submitted for statistical analyses.

Windowed cross-correlation analysis was computed at 50 ms lag times each way in 5 ms bins, using spiking data from the 0.8 mA PH stimulation epoch from the intensity step protocol. In the temporal domain, 50 ms window with 90% overlap were used. Cross-correlation of firing rates using the 250 ms segment during the stimulation was also computed between structures recorded.

### Histology

Upon the completion of data collection, rats were transcardially perfused with saline, followed by 10% formalin solutions. Their brains were extracted and kept at 4°C in cold 10% formalin solution for at least 24 hrs before the addition of 30% sucrose solution in preparation for cryostat sectioning. Brains were then sliced at 35 μm and mounted onto glass slides. Digital photographs of electrode tracks were taken for subsequent histological verification of recording areas. The recording sites were determined through estimation of the deepest point of the track, and notes regarding the depth of the electrode made during each experiment. Pontamine dots created by 50 μA constant current application from glass electrode recordings also supplemented the reconstruction. Only data from those units that were recorded within and immediately at the anatomical boundaries of each structure were included for analyses.

### Statistical Analysis

Repeated measures analysis of variance (ANOVA) was computed using SPSS 15.0 (SPSS Inc., Chicago, IL). All statistics are reported with Greenhouse-Geisser correction, regardless if the assumption of equal variances was upheld. Polynomial contrasts were used to determine the relationship between spiking and responses from intensity and duration step protocols. All post-hoc comparisons were corrected with the Bonferroni method to determine their statistical significance, but uncorrected *p*-values are reported in the text.

## Results

### Spiking Properties

Cells recorded from all target areas generally fell within known firing patterns described previously. Many STR neurons were relatively silent, with some neurons firing from a single action potential roughly once a minute and some were completely silent, but spiking can be elicited by PH stimulation readily. For the rest of the nuclei of interest, persistent spiking at high rates was observed with very few displaying oscillatory behaviour linked to ongoing cortical UP-DOWN states. Those that do display phase-locking to cortical UP-DOWN states were mostly found in the STR or the STN (see Figure 1 for an example). In this study, no attempt was made to dissociate different cell types recorded in a given structure. The averaged firing rates from the target nuclei are summarized in Table 2.

**Figure 1.**
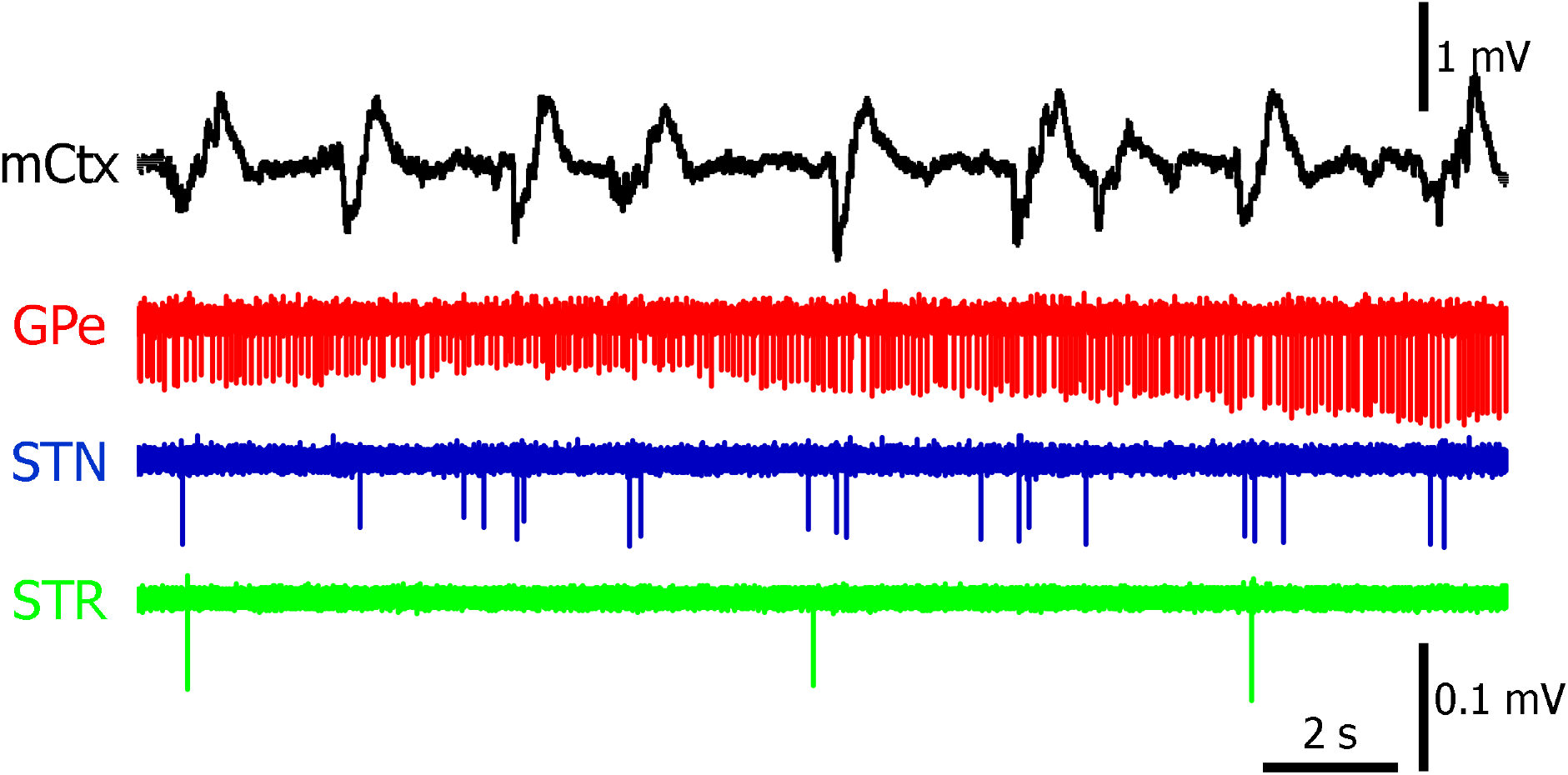
Raw data traces from an example of three concurrent basal ganglia single unit recordings with motor cortex LFP. Specifically, tonic, high-frequency firing can be seen in the GPe trace (red), whereas a more modest activity phase entrained to the cortical UP states can be seen in the STN (blue) and STR (green).

**Table 2.**
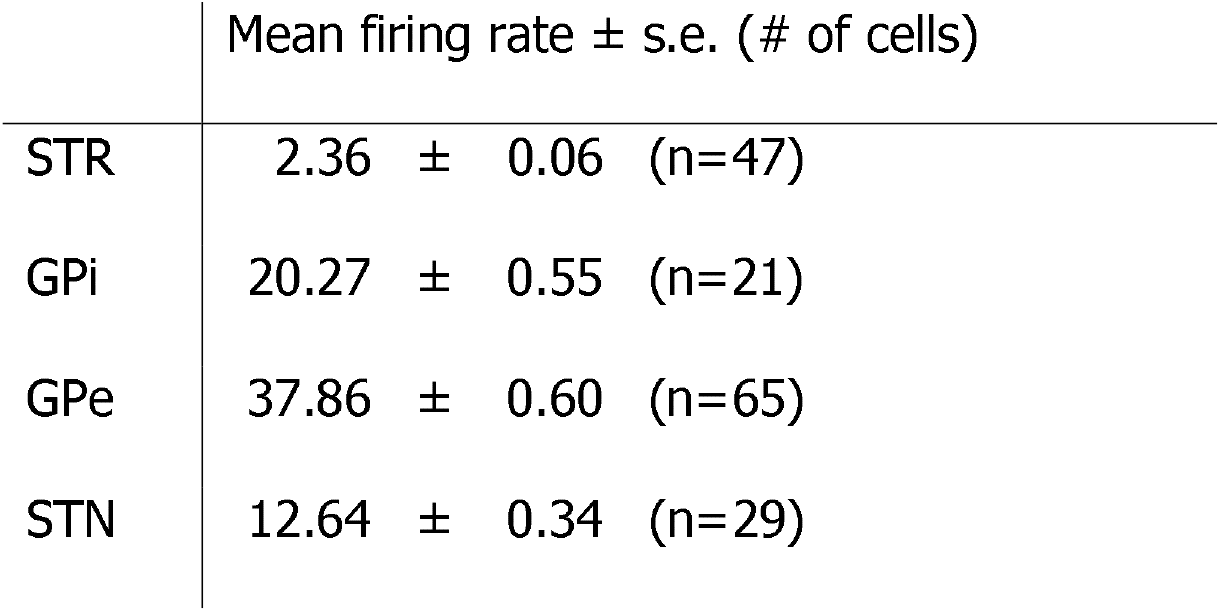
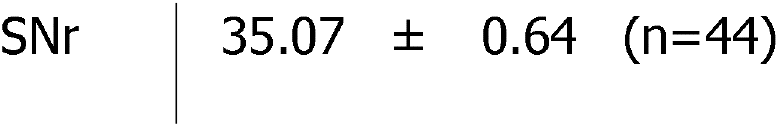
Mean firing rate and standard error of recorded basal ganglia structures.

### Basal Ganglia Response to PH Stimulation

Striatal neurons fired at low rates, if at all. The application of PH stimulation dramatically increased STR firing rate (Figure 2a, highlighted in red), which appeared to stabilize at a lower rate compared to the initial response to PH stimulation as a function of time (Figure 2a, highlighted in green). Elevated firing rates post-PH stimulation were also observed. The EP had a much higher basal firing rate compared to STR, but PH stimulation did not appear to have a discernable effect on the EP firing rate, despite a sustained increase of spike rate following the termination of PH stimulation (Figure 2b, bottom). Closer examination of the raster plot for EP suggests much heterogeneity exists in EP neurons in response to PH stimulation, in addition to the non-stationarity of individual neuronal firing rates during PH stimulation (Figure 2b, top). Recordings from the GPe reveal an initial excitatory effect at the onset of PH stimulation that fell back to baseline levels within ~1 s (Figure 2c, bottom). The firing rate or GPe appears to be at a slightly elevated level compared to baseline rates before the sustained spiking at even higher rates emerges at the end of the PH stimulation train. The raster plot suggests a relatively stationary response from the majority of cells during PH stimulation (Figure 2c, top). Similar to the changes described for GPe, neurons in the STN also maintained a slightly elevated firing rate at the duration of PH stimulation (Figure 2d, bottom). The same sustained increase in firing rate was also found after stimulus termination in the STN. The SNr is the only structure that displayed a tendency of decreasing its firing rate during PH stimulation, with the sustained increase of spiking observed again post-stimulus train (Figure 2e). In Figure 2f, recordings from two additional areas that have links to the basal ganglia are presented. Zona incerta (ZI) did not appear to respond to PH stimulation, with a slight decreasing trend of spiking until the termination of the stimulus train, where an elevated level of spiking was also observed. In the superior colliculus (SC), an observable decrease of spiking was observed initially, with more modest reductions of spiking for the rest of the stimulus train. Again, the same elevated firing rate can be seen at the termination of the stimulus.

**Figure 2.**
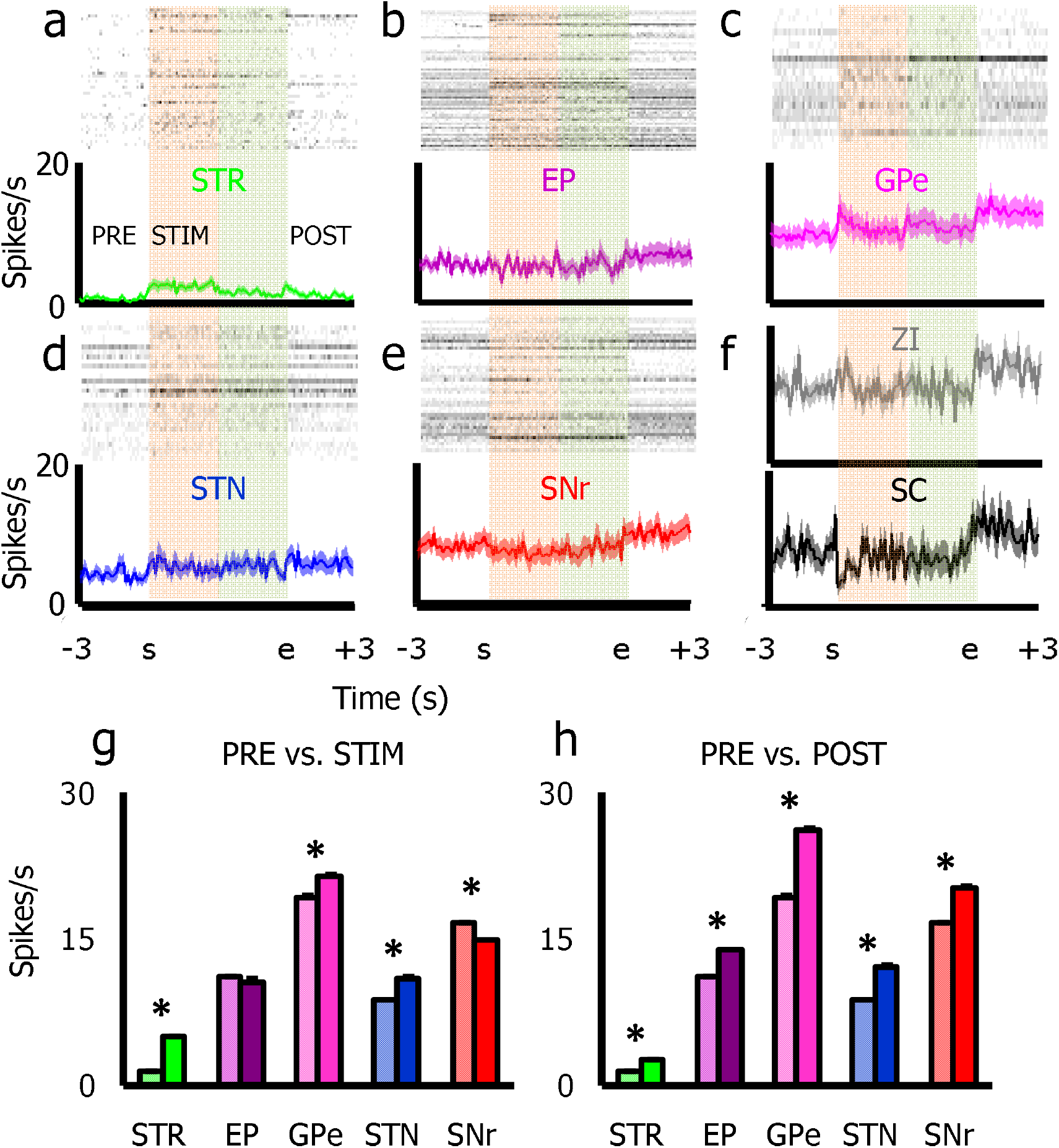
Spike rasters and averaged firing rates from two sets of PSTHs based on 10 s continuous PH stimulation. The first one consists of a 3 s pre-stimulation period (the first non-highlighted region), followed by the onset of PH stimulation (highlighted in red). This is joined to the last 3 s of PH stimulation (highlighted in green) and followed by another 3 s of a poststimulation period. (a) Ph stimulation increased the spiking rate of the STR during the stimulus train, which steadily decreased as the train continued, ending with elevated firing levels up to 3 s post-stimulation. (b) PH stimulation did not have any observable effects on EP activity, but elevated firing rates persisted post-stimulus. (c) In the GPe, PH stimulation appeared to transiently increase spiking, with firing rates just above the 3 s pre-stimulation baseline. Again, an elevated level of spiking was observed post-stimulus. (d) A modest increase in spike rate was observed in the STN, which remained relatively stable throughout the stimulus train, before further modest increases after the termination of it. (e) The only basal ganglia structure examined to show an inhibitory effect of PH stimulation in the SNr. The decreased spiking rate is relatively stationary during the PH stimulus train, and the post-stimulus increase in spiking rate was also observed despite the inhibitory response. (f) PSTHs constructed from data collected in the ZI and SC. Apart from a sharp, transient decrease in spiking from the SC, both spiked at levels comparable to baseline during PH stimulation. The post-stimulus increase in spiking rate was also observed in these two areas. (g) Differences in firing rates between an averaged 3 s baseline compared to the 3 s immediately after PH stimulation. The STR, GPe, and STN all demonstrated an increase in spiking, while there was a decrease in EP and SNr spiking activities. (h) Comparing the pre-stimulation and post-stimulation, 3 epochs revealed that all areas examined increased spiking activities after PH stimulation in all areas. * <.00005, see Table 3 for details.

Quantitative analyses reveal that changes in basal ganglia firing rates in response to PH stimulation described above are all statistically significant, except for the slight decrease detected in EP, which was not found to be statistically significant (Figure 2 g). The increase of spiking at the termination of the PH stimulation train was observed in all structures examined, and were all found to be statistically higher than baseline firing rates immediately prior to PH stimulation (Figure 2f). Summaries for individual statistics are presented in Table 3.

**Table 3.**
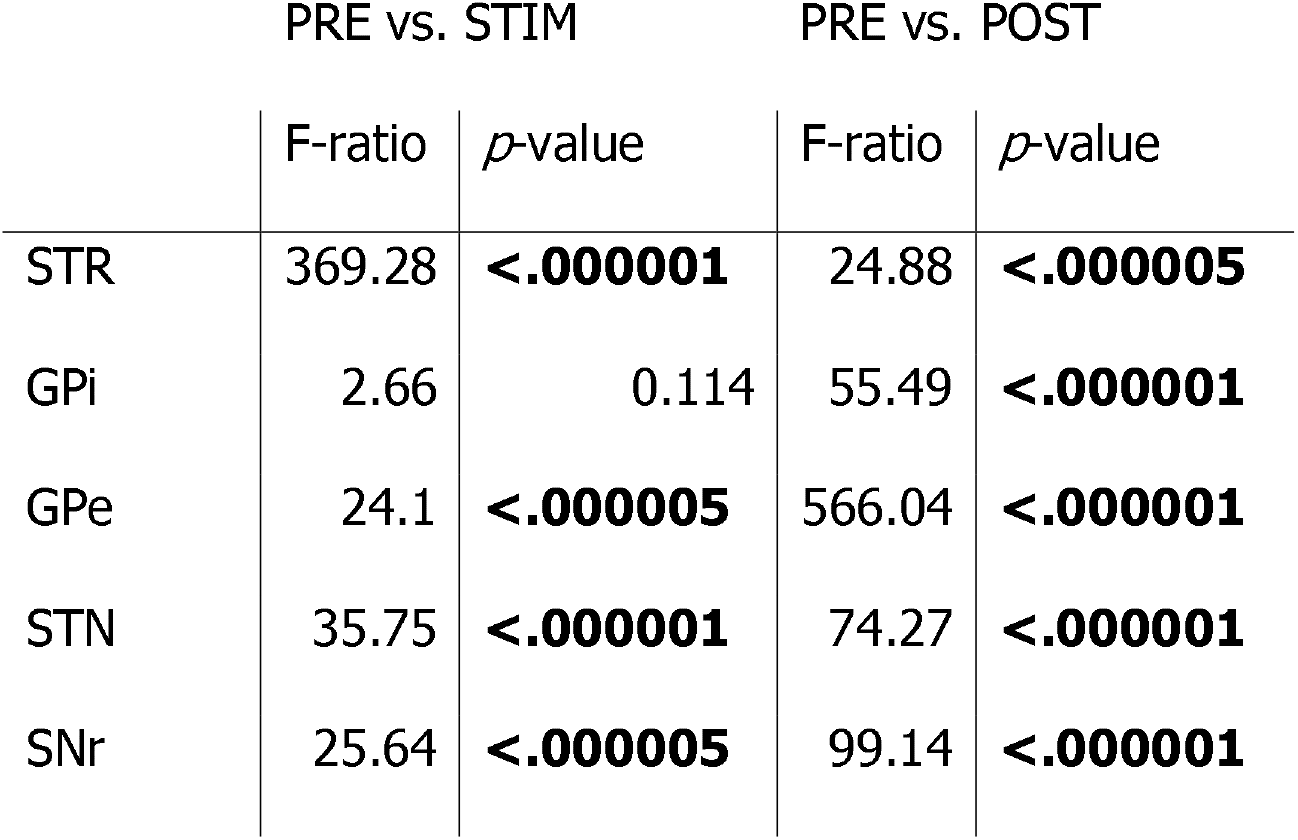
Firing rate changes in response to PH stimulation.

### The Effects of PH Stimulation Intensity on Basal Ganglia Responses

The effects of increasing PH stimulation intensity were varied between examined basal ganglia nuclei. There was a robust intensity-dependent response from the STR (Figure 3a), where spike rate linearly increased with increasing intensity (F_(1,46)_= 4.34, *p*< .05), despite an increase of variance as a function of intensity. This was not the case in EP, where the slope of intensity-spike rate function was relatively flat (Figure 3b, insert). In contrast, the GPe showed a statistically significant trend for a linear relationship between spike rate and PH stimulation intensity (F_(1,64)_= 5.82, *p*< .05). Neither the STN (Figure 3d) nor SNr (Figure 3e) demonstrated any linear changes as a function of increasing PH stimulation intensity. However, both areas did show significantly higher order polynomial trends (cubic and 7^th^ order, respectively) in relation to PH stimulation intensity.

**Figure 3.**
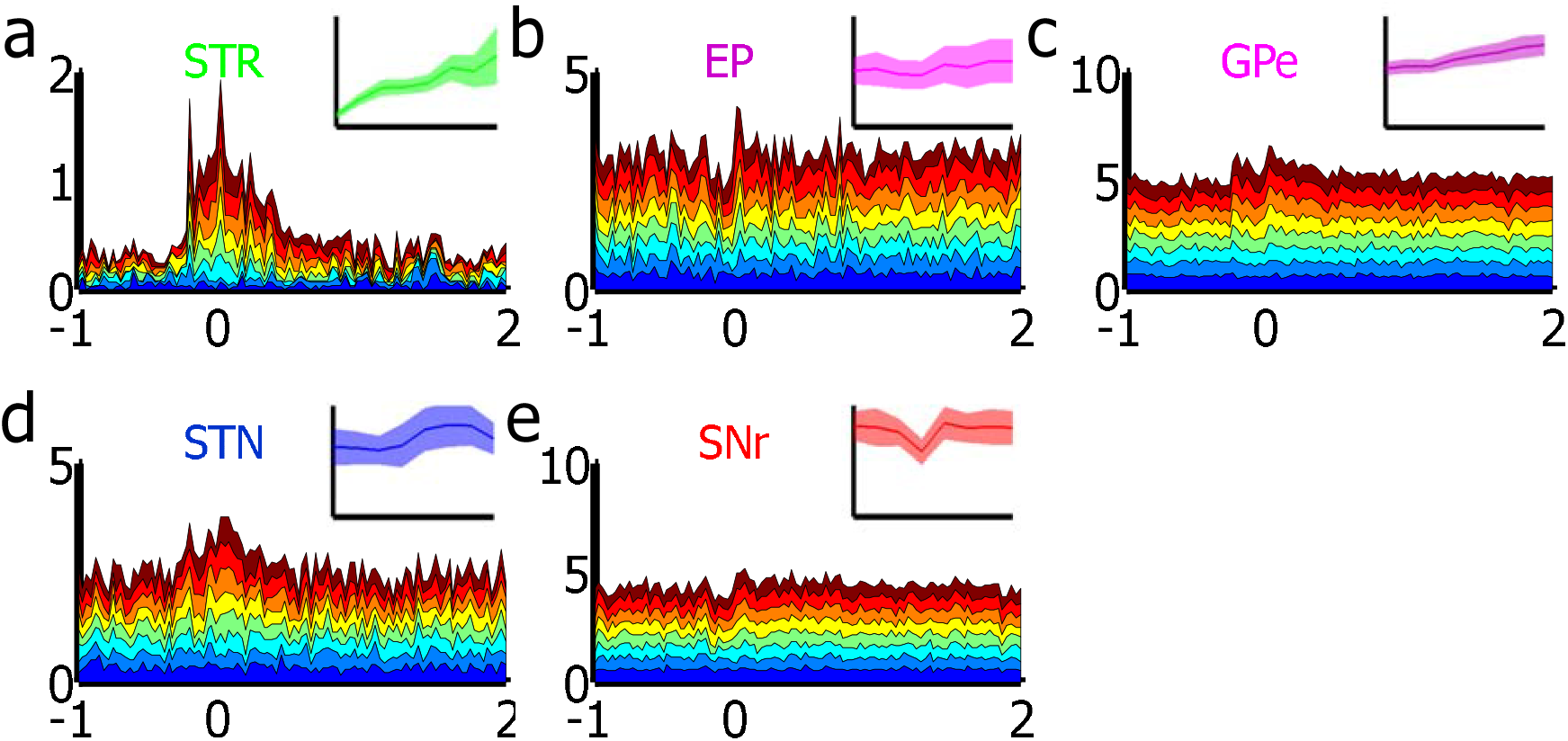
Basal ganglia responses to increased PH stimulation intensity. (a) In the STR, a linear relationship between spike rate and stimulation intensity was evidence (insert), with a clear demonstration of sustained post-stimulus increases in spiking. (b) There was no discernable effect of PH stimulation intensity on EP responses (insert), despite an apparent dip in firing rate associated with 0.2 and 0.4 mA steps. A rebound-like increase in spiking activity at the termination of PH stimulation was also observed. (c) More GPe spikes were evoked with increased stimulation intensity. A biphasic response similar to those described for longer trains of stimulation was seen in the GPe, with an initial transient in spiking rate, followed by a drop towards the baseline. The rebound-like transient increase in spiking after the PH stimulus train was also evident. (d) There was a cubic function associated with the STN spiking response to increased stimulation intensity (insert). A general increase of spiking is associated with the stimulus train. (e) There was no linear relationship between spiking rate and stimulation intensity in the SNr. In fact, the variances were so high between each intensity step that the only statistically significant trend was of the 7^th^ (highest possible) order. Similarly, brief inhibitory responses seen in the EP were detected at suprathreshold intensities of 0.3 and 0.4 mA. Rebound-like increases in spiking were also evident.

### The Effects of PH Stimulation Duration on Basal Ganglia Responses

Increasing stimulus duration linearly increased spiking rate in the STR (F_(1,46)_= 6.24, *p*< .05), despite some inconsistency at 200 ms (Figure 4a, insert). None of the other structures demonstrated any statistically significant relationships between spiking rate and increased PH stimulation duration (Figures 4b-e). The responses from these other areas were generally the same between different duration steps, and the responses were relatively more variable, as indicated by the amount of variance in spike rate in each nucleus.

**Figure 4.**
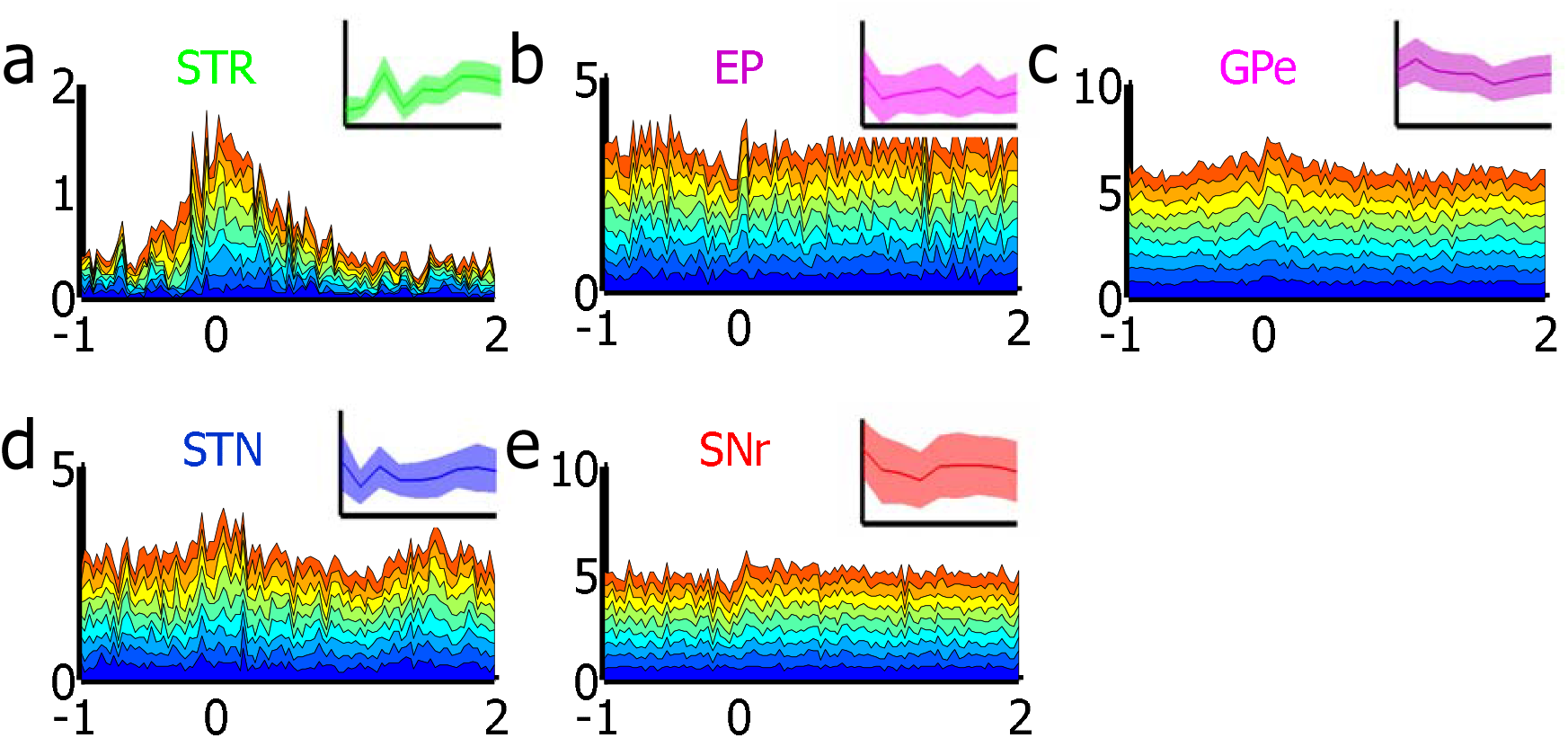
Basal ganglia responses to increasing PH stimulation duration. (a) Despite of the initial sharp increase of spike rate at suprathreshold levels, the steady increase of STR spiking rate is a function of increased stimulus duration. (b) Apart from having the time-course similar to those described in the intensity step, no relationship between stimulus train duration and spike rate was observed in the EP. (c) A decreasing trend was found between stimulus train duration and firing rates in the GPe, but was not statistically significant. (d) A flat slope in the stimulus train duration-spike rate plot suggests a lack of coupling between the two variables in the STN. (e) As mentioned in the previous figure, much variation exists within the SNr in response to PH stimulation. In addition to large variances, the lack of any observable trends of averaged responses across duration steps suggests no effect of PH stimulation duration changes on the SNr spiking rate.

### Correlated Responses to PH Stimulation between the Basal Ganglia

The correlated spiking changes between the basal ganglia nuclei examined in response to PH stimulation were investigated by using data obtained from the 0.8 mA stimulus train from the intensity-step protocols. In Figure 5, the top right half of the triangular matrix contains the windowed cross-correleogram of 0.8 mA stimulus train epoch that spanned from 750 ms before stimulation onset and the 2.25 s following it. Warmer colours represent the magnitude of positive correlation coefficients and cooler colours represent the magnitude of negative correlation coefficients. The plots clearly indicate an evoked response from all examined areas, with the exception of SNr. The time lags are calculated from 5 ms bins, therefore the striated distribution seen during the stimulation in every second bin represents the periods when stimulation artifacts preclude spiking activity. This pattern is more evident in EP and GPe, two nuclei that demonstrate high tonic firing rates. The same pattern is also evident in the STN, virtually absent from the SNr. The STR also demonstrates this pattern to a lesser extent, a result of interactions between increased, albeit at a relatively lower spiking frequency leading to fewer overall spikes lost to each stimulation artifact, in turn leading to higher auto-correlation coefficients.

**Figure 5.**
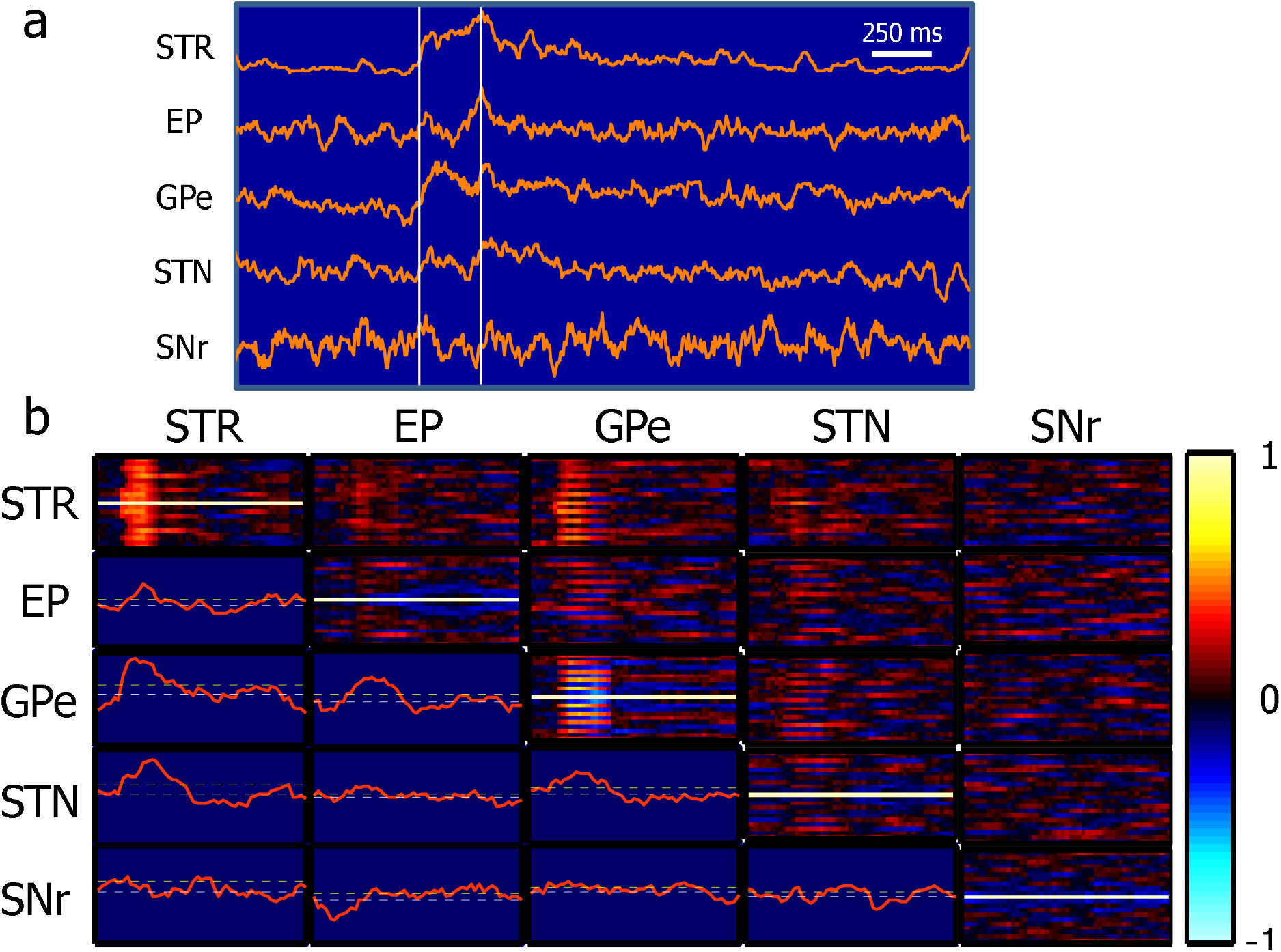
Cross-correlation analysis on basal ganglia spiking rates in response to 0.8 mA PH stimulation. (a) Averaged, normalized, and smoothed (5-point moving average of 5 ms bins) firing rate functions across the 3 s epoch of data consisting 750 ms lead in time to the stimulus train and 2 s following the cessation of the stimulation. While the stimulus increased STR firing rates, an inhibitory response was seen in the EP, a biphasic response for the GPe, and an apparent lack of change in the STN and SNr. The rebound-like increases in spiking after the stimulus can clearly be seen in GPe and STN responses. (b) Time-cross-correlation representation (top right triangular matrix) and ordinary cross-correlation (bottom left triangular matrix) of 3 s data, including the 0.8 mA PH stimulation train used in intensity step protocols. The diagonal represents time-auto-correlation representations from each structure. These plots show the lags in ms on the y-axis, spanning 50 ms in each direction, and the x-axis spanning the full 3 s of the original data epoch. Generally, the apparent oscillatory changes of correlation coefficients found during the stimulation are most likely a result of stimulus artifacts saturating the amplifier at 100 Hz (10 ms intervals), obscuring spiking activity at relevant intervals. Firing rates of the STR showed most correlated changes with other basal ganglia nuclei, with the exception of the SNr, which shares no coupled changes in firing rates with other nuclei. See text for more details.

The bottom half of the triangular matrix represents the time-dependent cross-correlation function of paired structures at a lag time of zero. The STR shows strong activation by PH stimulation, as indicated by the auto-correleogram but shares different time-courses with other structures of time-dependent spike rate. In the same column, correlated firing rate changes between STR and EP appear to be low, with a slower latency to reach the peak at the point close to the cessation of the stimulus train. In contrast, STR-GPe firing rate changes share much higher correlations and peak almost immediately after stimulus train onset. There was a somewhat biphasic response when STR firing rates were correlated with STN firing rates, with an initial rapid increase at stimulus train onset and further increased towards the end of the train. There did not appear to be any correlated firing rate changes between the STR and the SNr. The EP shared the highest correlated firing rates with the GPe towards the end of the stimulus train and maintained similar changes in firing rates briefly after stimulus cessation. There was a very modest increase in the correlated firing rate between EP and STN, while there is an indication that SNr had changes in firing rates that were in the opposite direction compared to EP, and PH stimulation appeared to normalize it. GPe demonstrated modest correlated firing rate changes with the STN, demonstrating a fall of correlation coefficients towards baseline after the stimulus train had terminated. The rest of the plots depicting SNr firing rate changes in relation to the STN and GPe revealed no significant interactions.

## Discussion

In this study, the response of five basal ganglia nuclei to PH stimulation was investigated. The data suggest PH stimulation has an overall excitatory effect on the basal ganglia, as evidenced by post-stimulus increases in spiking rate in all nuclei examined. Clear stimulus-bound increases of spiking rate as a function of stimulus intensity were detected in the STR and GPe, suggesting PH exerts more control on the excitability of these two nuclei. Changing the duration of the PH stimuli did not yield any effects other than increasing spiking in the STR. Finally, by correlating changes in firing rate in response to a short, high-intensity stimulus train, it was shown that the STR shows correlated firing changes with practically all other basal ganglia nuclei examined, with the exception of SNr, which demonstrated no correlated changes of spiking rate with any other basal ganglia nuclei.

One of the most robust effects demonstrated in this study was the high fidelity of PH stimulation eliciting an excitatory response in the STR. Although there are some directed projections from the PH to relatively localized areas in the PH, it is unlikely that these direct projections are the main determinant of the observed response. Instead, previous works have shown that PH stimulation results in global cortical activation, evoking cortical desynchronization and increased spiking activities (Young et al., 2011; Young and Bland, 2021), which may serve as one of the polysynaptic pathways PH may mediate its excitatory effects on the STR. Alternatively, previous work has also shown the PH-thalamic projections can drive thalamic, and subsequently cortical activity (Young et al., 2011; Young and Bland, 2021). Particularly, modest amounts numbers of PH fibres found in the ventrolateral thalamic nucleus were suggested as the neuroanatomical substrate for the powerful and spatially selective excitatory effects observed. The PH is known to project densely to the parafascicular and centromedian (intralaminar) nucleus of the thalamus (Veazey et al., 1982; Vertes et al., 1995), of which both provide thalamic innervations into the STR (Mengual et al., 1999; Van der Werf et al., 2002). Given its dense innervation to the intralaminar nuclei and excitatory effects on a wide range of cortical areas, it is likely the robust STR effect observed here is imposed by multiple polysynaptic pathways from the PH, through the thalamus and the cortex to elicit increased STR neural excitability. Striatal neurons are notoriously difficult to activate (even more so under anaesthesia), most likely requiring both cortical and thalamic glutamatergic inputs to arrive in a precise temporal window to be effective (Wilson et al., 1983; Wilson, 1986). The observation that some silent neurons recorded within the STR fire exclusively in response to PH stimulation also supports the notion that converging excitation to the STR principal cells is necessary to elicit action potentials.

The other consistent effect reported here is the elevated level of activity at the termination of the PH stimulus train, regardless of initial and/or sustained responses to PH stimulation itself. This effect was observed in all examined nuclei, suggesting it is not an effect driven by specific extrinsic inputs (such as the cortex) and propagated within the basal ganglia, since powerful STR excitation should result in inhibition of EP, GPe, and SNr, which would also result in the inhibition of the STN or demonstrate a more integrative response as those described during UP-DOWN states in the anaesthetized rat (Magill et al., 2004a; Walters et al., 2007). However, STN excitation can be driven indirectly by the cortex (Magill et al., 2004b) during and after PH stimulation, which may subsequently drive excitatory responses in all other structures examined here, of which all receive STN afferents (Deniau et al., 1978; Nauta and Cole, 1978; Van Der Kooy and Hattori, 1980; Carpenter et al., 1981; Kita et al., 1983; Kita and Kitai, 1987; Parent and Hazrati, 1995b; Plenz and Kital, 1999; Sato et al., 2000). It is not possible to employ a solely cortical-driven mechanism to explain an overall excitatory effect in all basal ganglia nuclei examined here with the available data. A more parsimonious explanation invoking the undeniable role of the PH has in general arousal may be sufficient to account for the changes. Apart from its hypocretin/orexin positive neurons (Peyron et al., 1998; Nambu et al., 1999; Abrahamson and Moore, 2001; Cheng et al., 2003; Hahn, 2010), the PH also has direct connections to neuromodulator centres, such as the locus coeruleus, midbrain dopaminergic nuclei and the serotonergic raphe (Vertes et al., 1995; Vertes and Crane, 1996) to trigger an increased state of arousal, affecting the whole brain. The previous observation of sustained cortical and thalamic persistent tonic firing (Young et al., 2011; Young and Bland, 2021), as well as the same changes observed in the superior colliculus and zona incerta reported here, support the claim that in addition to its direct synaptic effects on downstream neurons, the ability for PH stimulation to globally evoke an increased state of arousal imposed on the rest of the brain may be the most salient feature of its activation.

The other interesting observation is the response of GPe to PH stimulation. GPe is the only nucleus other than STR that demonstrated a linear response of spike rate to increasing stimulation intensity. Since the GPe has virtually intrinsic connections with the rest of the basal ganglia, there are limited possibilities of how PH stimulation may mediate this robust effect. The main input to the GPe is from the STN (Parent and Hazrati, 1995b), which exerts powerful excitatory effects on the GPe (Smith and Parent, 1988; Robledo and Feger, 1990). Through increased cortical excitability and activity, increase spiking in the GPe can be mediated through the STN via the cortex. However, data reported here suggest a weak relationship between spiking rate and PH stimulation intensity in the STN, despite sharing correlated increases in spiking rate along with the STR and GPe. Particularly, the observed increase of spike rate in relation to GPe was not as prevalent as the relationship shown with the STR. Therefore, it is unclear if STN is involved in GPe activation during PH stimulation. The ambiguity is also partly due to the intimate STN-GPe functional connectivity (e.g. Plenz and Kital, 1999), which may result in nonlinear interactions between the two, yielding inconclusive results based on linear measures employed here. Alternatively, PH projects almost exclusively to the part of GPe targeted in the current study (Vertes et al., 1995). Therefore, the possibility exists that the robust excitatory effects of PH stimulation on GPe spiking activity are direct, which would also account for the ambivalent response from the STN. Arguing against the hypothesis regarding linearity of evoked responses solely on an anatomical basis, both the SC and ZI also receive innervations from the PH, with ZI being one of the more densely innervated areas by the PH (Vertes et al., 1995; Vertes and Crane, 1996). However, no increases in spiking rate can be seen from these two areas during PH stimulation. While the contamination of stimulus artifacts may have prevented the detection for such an increase, the GPe does fire at similar frequencies compared to ZI and SC, yet an increase of spiking was detected. Additional recordings sampling the whole extent of the GPe are required to verify the validity of this hypothesis.

The highly interconnected nature of the basal ganglia renders the interpretation of data collected from isolated recordings from different structures difficult. With this preliminary report, directed paired recordings can be made to test the hypothesis generated by current observations. For example, the relationship of GPe-STN responses to PH stimulation should be addressed by paired recordings to determine the direction of influence during PH stimulation to clarify the apparent lack of STN involvement in GPe excitation suggested by available data. The SNr is the only structure that demonstrates a statistically significant decrease of spiking with PH stimulation and shows little coupling of firing rate changes to all other basal ganglia structures recorded here. It is possible that in the absence of direct excitatory modulation from the STN, STR inputs predominate and result in inhibitory responses. Circuit analysis based on concurrent multi-site recordings would be beneficial to elucidate the effects of PH stimulation on the basal ganglia.

The major weakness of the present study is inferring changes in spiking in the presence of stimulus artifacts. Stimulus artifacts not only make spike sorting from MUA traces difficult due to the loss of signal-to-noise ratio due to lower electrode impedance, but it also obscures the spiking activity centered on the artifact itself. For neurons that have a slow spike rate, such as STR neurons reported here and thalamic and cortical neurons reported previously, it is possible to effectively detect robust increases in spiking rate even with stimulus artifacts present. However, for neurons that have higher firing frequencies, and therefore the likelihood of exhibiting a tonic firing pattern, stimulus artifacts would have a higher probability of obscuring action potentials or simply change the spike waveform due to the direct component associated with the relatively slow decay of the signal back to baseline from saturation. Therefore, although certain relationships were uncovered here regarding the effect of PH stimulation on basal ganglia firing rates, the only concrete conclusion that can be made is that the PH does not abolish neural activity in the basal ganglia circuitry. Recent developments of optically based stimulation with genetically engineered bacteriorhodopsins expressed in mammalian systems can overcome issues encountered with electrical stimulation (Boyden et al., 2005; Airan et al., 2007; Zhang et al., 2007).

This study reports the effects of PH stimulation on basal ganglia spiking activity in urethane anaesthetized rats. It was found that PH stimulation elicited strong stimulation-dependent excitatory responses from the STR and GPe. Basal ganglia responses during PH stimulation and immediately after it cannot be explained by activity propagation within the basal ganglia alone. Cortical and thalamic involvement is most likely necessary for the powerful effects of PH stimulation on STR neurons, while the robust effect of GPe activation may be mediated by direct PH projections in the absence of STN involvement. Stimulation of the PH undoubtedly triggered an increase in general arousal, as all the regions recorded to date demonstrated an increased number of spiking following the stimulation. Concurrent multisite recordings are necessary to elucidate the relative contribution of a common source (e.g. arousal systems) and specific circuit-dependent properties intrinsic to the basal ganglia to observations reported here.

## Acknowledgements

This research was supported by Natural Sciences and Engineering Research Council (NSERC) research grant to BHB (A9935). CKY was supported by Parkinson Society Canada (PSC) and Parkinson Society of Southern Alberta (PSSA).

## Contributions

CKY conceived the study, carried out all experiments and analyses. BHB provided supervision and secured funding. CKY prepared the manuscript and both CKY and BHB reviewed and edited the manuscript.

